# Effectors from a Bacterial Vector-Borne Pathogen Exhibit Diverse Subcellular Localization, Expression Profiles and Manipulation of Plant Defense

**DOI:** 10.1101/2021.09.10.459857

**Authors:** PA Reyes-Caldas, Jie Zhu, A Breakspear, SP Thapa, TY Toruño, L Perilla-Henao, C Casteel, C Faulkner, G Coaker

## Abstract

Climate change is predicted to increase the prevalence of vector borne disease due to expansion of insect populations. *Candidatus* Liberibacter solanacearum (*C*Lso) is a phloem-limited pathogen associated with multiple economically important diseases in Solanaceous crops. Little is known about the strategies and pathogenicity factors *C*Lso uses to colonize vector and host. We determined the *C*Lso effector repertoire by predicting SEC secreted proteins across four different *C*Lso haplotypes, investigated effector localization *in planta*, and profiled effector expression in vector and host. The localization of *C*Lso effectors in *Nicotiana* revealed diverse eukaryotic subcellular targets. The majority of tested effectors were unable to suppress plant immune responses, indicating they possess unique activities. Expression profiling in tomato and the psyllid *Bactericera cockerelli* indicated *C*Lso differentially interacts with its host and vector and can switch effector expression in response to the environment. This study reveals *C*Lso effectors possess complex expression patterns, target diverse host organelles and the majority are unable to suppress host immune responses. A mechanistic understanding of Lso effector function will reveal novel targets and provide insight into phloem biology.

## INTRODUCTION

Vector-borne diseases (VBDs) reduce agricultural productivity and disrupt ecosystems worldwide. Rising global temperatures are predicted to increase insect populations and fitness, exacerbating the dispersion of emergent VBDs (Deutsch *et al*., 2018). Some of the most widespread and devastating VBDs are associated with Liberibacter. Liberibacters are gram-negative, obligate, phloem-limited bacterial pathogens that are transmitted by different psyllid vectors onto plant hosts (Perilla-Henao & Casteel, 2016). Huanglongbing (HLB) disease, also known as Citrus Greening, is primarily associated with *Candidatus* Liberibacter asiaticus (*C*Las) and is considered the most important citrus disease worldwide (Singerman & Rogers, 2020). *C.* Liberibacter solanacearum (*C*Lso) is associated with economically important diseases on a variety of Solanaceous and Apiaceous hosts, including tomato (Tomato Psyllid Yellows) and potato (Zebra Chip).

Piercing-sucking insects, such as psyllids, insert their stylet into plant hosts and probe the apoplast as well as other cell types before reaching phloem. During probing, insects secrete watery saliva containing HAMPs (herbivore associated molecular patterns), and other pathogen-associated molecular patterns (PAMPs) derived from bacteria present in their body/salivary glands (Chaudhary *et al*., 2014; Jaouannet *et al*., 2014). Phloem responses to biotic stress or insect damage include the accumulation of nitric oxide (NO), reactive oxygen species (ROS) and Ca^2+^ in phloem bundles which are required for rapid systemic signaling. NO and ROS are required for the establishment of systemic acquired resistance while a calcium-ROS auto-propagation wave interacts with electric signals for induction of systemic wound responses (Gaupels *et al*., 2016, 2017). ROS and Ca^2+^ influx in vascular bundles leads to activation of occlusion proteins, callose deposition and phytoalexin accumulation (Huang *et al*., 2020).

Plant pathogens rely on secretion of proteins, called effectors, that modify their host. Liberibacter species lack specialized secretion systems for specific delivery of effector proteins into host cells, but harbor all the essential components of the general secretion (SEC) machinery for delivery to the bacterial periplasm. Liberibacter SEC effectors are secreted when using *Escherichia coli* as a surrogate (Prasad *et al*., 2016). Noncanonical secretion of periplasmic Liberibacter proteins likely occurs through outer membrane vesicles (OMVs), as OMVs have been well-characterized as an alternative route for secretion from the bacterial periplasm and have been visualized in *C*Lso (Nissinen *et al*., 2014; Magali *et al*., 2015; Jonca *et al*., 2021). Noncanonical effector secretion was demonstrated using heterologous expression of two *C*Las proteins (CLIBASIA_RS0045 and SC2_gp095) in the culturable surrogate *Liberibacter crescens* (Jain *et al*., 2015, 2018, 2019). Predicted *C*Las effectors are small, which predicts their ability to move symplastically *in planta* away from the sieve elements through plasmodesmata. SDE1, a small secreted *C*Las effector protein, moves far from the point of infection either by phloem transport or cell-to-cell movement through plasmodesmata (Pagliaccia *et al*., 2017). Phytoplasmas, another class of insect-vectored phloem-limited bacterial pathogens, also secrete effectors that move through plasmodesmata to manipulate their host (Tomkins *et al*., 2018).

To date, only a handful of Liberibacter effectors from any species have been characterized. *C*Las Sec-dependent effector 1 (SDE1) is preferentially expressed in citrus and periwinkle hosts, suppresses plant defense by targeting plant immune proteases, and transgenic plants expressing SDE1 phenocopy leaf blotchy mottle disease symptoms (Pagliaccia *et al*., 2017; Clark *et al*., 2018, 2020). *C*Las SDE15 and *C*Lso-hypothetical protein effector 1 (HPE1) were also reported to suppress plant cell death (Levy *et al*., 2019; Pang *et al*., 2020). Phytoplasma effectors such as TENGU and SAP11 target transcription factors involved in the production of phytohormones, inducing disease phenotypes such as witches broom and flower sterility in addition to impairing plant defenses and vector performance (Sugio *et al*., 2011; Tan *et al*., 2016; Dermastia, 2019). Despite the importance of bacterial effectors in promoting vector-borne disease development, little is known about the identity, conservation and role of *C*Lso effectors. Breakthroughs in understanding Liberibacter pathogenicity mechanisms could be achieved using *C*Lso as a model system due to the ability to study disease in *Nicotiana tabacum, Nicotiana benthamiana* and *Solanum lycopersicum* (tomato) (Huang *et al*., 2021).

*C*Lso is the most recently evolved species within the Liberibacter genus and is divided into 12 haplotypes with different vector and host preferences; three are associated with diseases in Solanaceous plants (A, B and F) and six are reported in a wide range of Apiaceous crops (C, D, E, H, Cras1 and Cras2) (Sumner-Kalkun *et al*., 2020; Thapa *et al*., 2020). *C*Lso haplotypes cause different symptomatology (Mendoza-Herrera *et al*., 2018). *C*Lso haplotypes A and B are capable of systemically colonizing and propagating in both vector (*Bactericera cockerelli*) and host (tomato and potato). Because of their inability to be cultured and their specific phloem localization, understanding the etiology and biology of the diseases caused by Liberibacters has been challenging (Huang *et al*., 2020). Currently there is no genetic resistance in cultivated plant germplasm and increasing resistance to insecticides pose a high risk for disease epidemics (Chávez *et al*., 2015; Szczepaniec *et al*., 2019).

Here, we studied *C*Lso effectors to gain insight into disease development. SEC secreted effectors were identified from four different *C*Lso haplotypes. *C*Lso effector localization was analyzed after transient expression in *Nicotiana*. We evaluated the ability of *C*Lso effectors to suppress early markers of plant defense in response to perception of pathogen features. Despite diverse subcellular localizations, the majority of tested effectors were unable to suppress host immune responses, indicating they possess unique activities. Expression profiling in tomato and the psyllid *B. cockerelli* highlighted three patterns of effector expression: early and late acting effectors, preferential expression in the vector or host, and constitutive effector expression across vector and host. This study highlights promising *C*Lso effectors for future investigation.

## MATERIALS AND METHODS

### Plant Materials and Growing Conditions

*Nicotiana benthamiana* wildtype and the transgenic line SRLJ15 expressing Aequorin (Segonzac *et al*., 2011) plants were grown in a controlled environment chamber at 26°C and 12h light/dark photoperiod with light intensity of 180 µE m^−2^ s^−1^. Tomato plants (*Solanum lycopersicum* cv Money Maker) were grown under controlled conditions at 26°C and 12h light/day photoperiod. *Bactericera cockerelli* psyllids carrying *C*Lso haplotype B of the Western biotype were reared in tomato plants inside a 60□×□60□×□60-cm Bugdorm insect rearing cage (BioQuip Products, Rancho Dominguez, CA). All experiments were performed using synchronized colonies. To synchronize *B. cockerelli* colonies, 15-20 adult psyllids were collected from infected symptomatic or clean plants and transferred to healthy 5-week old tomato plants inside a 24.5□×□24.5□×□63-cm Bugdorm insect rearing cage (BioQuip Products, Rancho Dominguez, CA). 72h post-infestation, the adult psyllids were removed, and the plants were kept 21-26 days until new adults emerged.

### Effector Prediction

*C*Lso genomes for haplotype A (JQIG01,JMTK01, JNVH01), B (CP002371.1) C (LVWB01, LVWE01), and D (LLVZ01) were used to predict secreted proteins with a signal peptide (SP), using SignalP v3.0 and SignalP v4.1 (Bendtsen *et al*., 2004; Petersen *et al*., 2011). Proteins larger than 35KDa and containing predicted transmembrane (TM) domains (TMHMM v2.0, Krogh et al., 2001) were removed. Effector conservation analyses were performed in pairwise comparisons using BLASTP and BLASTN (Gish *et al*., 1990). The threshold of percent similarity for presence/absence was 40%. The presence of eukaryotic nuclear localization signals was predicted using LOCALIZER web server (Sperschneider *et al*., 2017). All predicted effectors are listed in Table S1.

### Phylogenetic analysis

Orthologous genes of *C*Lso isolates were predicted using the OrthoMCL v. 2.0 pipeline (Li *et al*., 2003). All-versus-all BLASTN comparison of all gene sequences for each species was performed, and orthologous genes were clustered by OrthoMCL v. 2.0. Multiple alignments of gene sequences were performed with PRANK v. 170,427 (Löytynoja, 2014). All the alignments were concatenated by FASconCAT v. 1.1, yielding a gene supermatrix (Kück & Meusemann, 2010). A maximum-likelihood approach was used to reconstruct the phylogenetic tree using RAxML v. 8.2 software (Stamatakis, 2014). The bootstrap was performed with 1,000 replicates. The resulting tree was visualized using FigTree v. 1.4.3 (Rambaut, 2012).

### Effector Cloning

To create N-terminal fusions with tGFP or C-terminal fusions with eGFP, we used the Golden Gate Toolkit for plants (Engler *et al*., 2014a). *C*Lso mature effectors (without signal peptide) were amplified from haplotype A and B infected tomatoes genomic DNA under standard PCR conditions using iProof (Biorad, Supplementary Table 1). Annealing temperatures of primer pairs were calculated using FastPCR (Kalendar *et al*., 2011). Amplicons of the expected size were purified and subcloned in the respective level zero acceptor for CDS1ns or CDS2 parts (Supplementary Table 3). Effector clones were confirmed by PCR and sequencing. To generate the fluorophore tagged fusions, confirmed effectors were cloned in the Golden Gate level 2 acceptor plasmid pICH86966 using a restriction-ligation reaction. To assess symplastic mobility of *C*Lso effectors, a new Golden Gate vector (FP08018-BsaI) was engineered. The vector contains a constitutively expressed, cell-to-cell immobile, nuclear localized tdTomato (transformation control) as well as a 35S promoter-*lacZα*-eGFP cassette. The *lacZα* is flanked by *Bsa*I recognition sites allowing PCR-amplified *C*Lso effectors to be introduced with a Golden Gate reaction. The vector was produced using Golden Gate assembly of pre-existing basic parts (Engler *et al*., 2014b) and newly created level 0 modules for *lacZα* and tdTomato. Control constructs containing nuclear localized tdTomato and a mobile eGFP (FP08024) or immobile 2XeGFP (FP08027) were also produced using Golden Gate assembly. Effector clones were confirmed by PCR, restriction digestion and sequencing. Final constructs were transformed in *Agrobacterium tumefaciens* GV3101. HPE19 was subcloned in the entry vector pENTR/SD/D-TOPO, following manufacturer instructions and cloned into the destination vector pTA7001(Li *et al*., 2013) using a Gateway LR reaction (Invitrogen). Sequences of destination vector inserts were confirmed and then transformed in *Agrobacterium tumefaciens* GV3101. All primers used for cloning are listed in Supplemental Table 2, plasmids are listed in Supplemental Table 3, and effectors are listed in Supplemental Table 1.

### Western blotting

To evaluate *C*Lso effector expression, individual effectors were cloned at tGFP fusions and expressed in *N. benthamiana* using *Agrobacterium*-mediated transient expression. *A. tumefaciens* GV3101 carrying effectors was induced with 100 µM acetosyringone for 1h and infiltrated at an OD_600_ = 0.5. Total protein was isolated from *N. benthamiana* leaves 24h post-infiltration (hpi) by grinding in 2X Laemmli buffer (Laemmli, 1970). Protein samples were separated by SDS–PAGE and immunoblotting was conducted using anti-tGFP at a concentration of 1:10000 (Invitrogen), followed by secondary anti-rabbit-HRP at a concentration of 1:3000 (Biorad). Positive signals were detected via chemiluminescence using the Super Signal West Femto Chemiluminescent Substrate (Pierce) and visualized using the Bio-Rad Chemidoc system.

### Confocal microscopy and effector mobility

*A. tumefaciens* GV3101 carrying effectors was induced with 100 µM acetosyringone for 1h and infiltrated at an OD_600_ = 0.5 into *N. benthamiana*. Imaging was performed at 24 hpi. Plasmolysis was performed using 1 M NaCl. Specific mCherry markers for the plasma membrane (pm-rkCD3-1007), ER (ER-rk CD3-959), Golgi (G-rkCD3-967) and peroxisome (px-rk CD3-983) were mixed in a 1:1 ratio with *C*Lso tagged effectors, co-infiltrated in *N. benthamiana* and visualized at 48hpi. All confocal microscopy was performed using a Leica SP8 confocal scope equipped with a 63X water objective. tGFP was excited at 488 nm, and the emission was gathered at 500 to 550 nm. mCherry was excited at 550nm and the emission gathered at 570 to 600nm. The chloroplast autofluorescence emission was gathered at 650 to 750 nm. Images were analyzed using the Leica Application Suite X (LAS X) software, 3D reconstructions were generated using the 3D visualization and analyses tools in the LAS X Core module.

For effector mobility assays, *A. tumefaciens* GV3101 carrying HPE1-eGFP, HPE8-eGFP, pF08024 (eGFP) and pF08027 (2xeGFP, Supplemental Table 3) were infiltrated at an OD_600_ = 0.0005. Imaging was performed at 48 hpi using a Leica SP8 confocal scope. eGFP was excited at 488 nm, and the emission was gathered at 495 to 540 nm. tdTomato was excited at 552 nm, and the emission was gathered at 561-616nm. Single transformation events were observed with a 20X objective in a Leica SP8 confocal scope.

### Trypan Blue staining and Electrolyte Leakage Assay

*N. benthamiana* leaves were infiltrated with *A. tumefaciens* GV3101 carrying either t-GFP, tGFP-HPE19 or BAX under as described above. 24 hours later leaves were infiltrated with 2μM dexamethasone for GAL4 promoter constructs. Leaves were detached 5 days post *Agrobacterium*-mediated transient expression and subjected to Trypan Blue staining and distained with Chloral hydrate (1.25g/mL) as previously described(McDowell *et al*., 2011). Electrolyte leakage was measured as previously described (Bolus *et al*., 2019). Samples from the different treatments were collected using a N°9 cork borer (0,79cm^2^) at 12h post Dex induction. Individual leaf discs were placed in a 12 -well tissue culture plate with 5mL of distilled water for 30 minutes. Water was replaced with 2mL of distilled water and leaf discs were kept at room temperature under constant 30µmols^-1^m^-2^ light. Conductivity (µS/cm) was measured using a Model 3200 conductance instrument (YSI) every 24 hours for five days. Statistical differences were detected using a repeated measures ANOVA and different groups were assigned using Tukey’s post-hoc test.

### ROS burst and cytosolic calcium accumulation assay

*N. benthamiana* wildtype at the two leaf stage was inoculated with *Agrobacterium* strain GV3010 at an OD_600_ = 0.5 containing the mature effector side by side with the control *Agrobacterium* strain GV3010 containing an empty vector (EV). For ROS, leaf discs were excised with a N°1 cork borer (0,48cm^2^) at 24 hpi and incubated overnight in water. Before measurement, the water was removed and 100 μL of assay solution (17 mM luminol, 1 μM horseradish peroxidase, and 100 nM flg22 [u1959ek130 Genescript] or 100 μg mL^−1^ chitin hexamers(O-ch16 Megazyme) was added to each well. Transient increase of cytosolic Ca^2+^ concentration was monitored in the *N. benthamiana* SLJR15 line (Segonzac *et al*., 2011). Leaf discs were excised at 24 hpi and incubated overnight in 1 μM native coelenterazin (Sigma). Before measurement, the solution was removed and 100 μL of assay solution (100 nM flg22 or 100 μg mL^−1^ chitin) was added to each well. Luminescence was measured using a TriStar Luminometer. Experiments included three biological replicates (individual plants) and were repeated three times (n = 9 plants). Statistical differences in ROS production or calcium accumulation were detected using multiple Mann-Whitney tests. Adjusted P values from the Holm-Sidak method were used to compute adjusted P values. *p<0.05, **p<0.005, ***p<0.0005, **\*\*\*\*** p<0.00005. Outliers were detected by the ROUT method and removed from the analyses.

### Time-Course Expression Analyses of *C*Lso effectors in tomato and psyllid

Three-week-old tomato *Solanum lycopersicum* cv Money Maker were infected under controlled conditions with fifteen newly emerged psyllids from a synchronized *C*Lso haplotype B colony. Psyllids were confined in a muslin bag, removed after 72h, flash-frozen in liquid nitrogen and stored at -80°C until RNA extraction. The leaves were inspected and eggs (if present) were carefully removed using tape. Midvein samples (∼50mg) were collected from the originally inoculated leaves at one or four-weeks post vector transmission. Six individual plants were included per experimental timepoint. Fifteen recovered psyllids were pooled for RNA extraction per infected plant. Total RNA was isolated using the TRIzol reagent (Invitrogen) following the manufacturer’s instructions. To guarantee DNA removal, an additional cleaning step with acid phenol was added before precipitation of the RNA by adding an equal volume of Acid-Phenol:Chloroform 5:1 solution pH 4.5 (Ambion). Total RNA was quantified with a Nanodrop 2000 spectrophotometer (Thermo Scientific). The *C*Lso titer for each sample was assessed using the RecA primers (Ibanez *et al*., 2014) and the three samples with the best Ct values for each timepoint were selected for further analysis. iTaq Universal SyberGreen One-Step RT-qPCR kit (Biorad) was used to perform RT-qPCR following the manufacturer’s protocol. 1µL of a 1:2 dilution of RNA (∼100-250ng) was used per reaction. Effector expression was quantified on the CFX96 real-time PCR detection system (Bio-Rad). Gene expression was calculated using the ΔCT method and normalized against the *GlnA* housekeeping gene. Primers were designed with Primer3 (Untergasser *et al*., 2012) and provided in Supplementary Table 4. Pheatmap R package was used for plotting the histograms and performing hierarchical clustering (Kolde, 2012).

## RESULTS

### 2.1 The *C*Lso SEC-effector repertoire varies in size and composition

To begin investigating how *C*Lso is able to colonize two distinct organisms and thrive in the phloem environment we predicted suites of SEC-dependent effectors across four haplotypes (A-D). *C*Lso genomes were downloaded from NCBI, including three from haplotype A (LsoNZ1, HenneA, R1), two from haplotype C (FIN111 and FIN114) and one from haplotypes B and D (CLso-ZC1 and ISR100, respectively; Fig 1a)(Lin *et al*., 2011; Zheng *et al*., 2014; Thompson *et al*., 2015; Wang *et al*., 2017). Genome CP002371.1 (CLso-ZC1) is completely finished, while the others are draft. SEC dependent effectors were predicted using SignalP and filtered to remove proteins larger than 35KDa as well as those with predicted transmembrane domains. We found 13 effectors that are conserved in all the haplotypes screened (core) and 27 that were found in several but not all haplotypes (variable). Each haplotype carried between 27-36 predicted effectors (Table S1), with haplotypes C-D having the smallest (27-29) and haplotypes A-B having the largest (34-36) repertoires (Fig 1b). In total, we estimate the repertoire of effectors across *C*Lso haplotypes to be ∼27, comprising 13 core, 6 variable, and 8 unique effectors (Fig 1b). These estimates could change with the sequencing of new strains and haplotypes.

**Figure 1.**
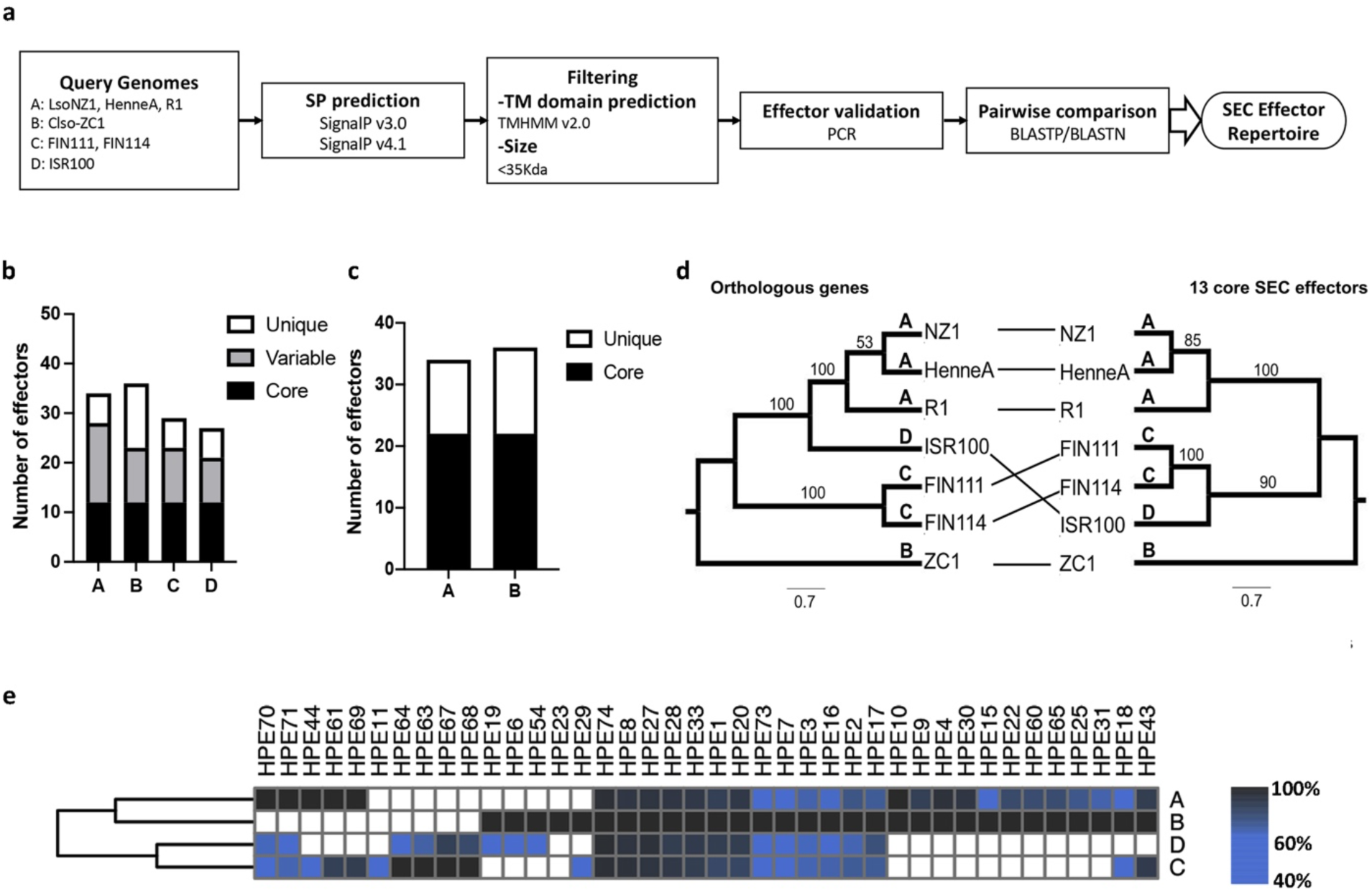
Prediction and classification of *C*Lso SEC-dependent effectors. **a.** Pipeline used to predict the *C*Lso effector repertoire. *C*Lso genomes for haplotype A (JQIG01, JMTK01, JNVH01), B (CP002371.1) C (LVWB01, LVWE01), and D (LLVZ01) were used to predict secreted proteins with a signal peptide (SP). Proteins larger than 35KDa and containing predicted transmembrane (TM) domains were removed. Effectors were PCR validated in haplotype A (LsoNZ1) and B (CLso_NZ1). **b.** *C*Lso effectors were classified into three categories: those present in all haplotypes (core), those shared only with one or two haplotypes (variable) and those unique to each haplotype. **c.** Effector repertoire comparison across multiple strains of haplotypes A and B. The criteria and genomes utilized are the same as in B. **d**. Left: *C*Lso phylogeny based on 815 orthologous genes. Right: *C*Lso phylogeny based on the 13 core effectors. A maximum likelihood approach was used to generate each phylogeny with 1000 bootstrap replicates. Bootstrap values are indicated at each node. **e.** Presence/Absence hierarchical clustered heatmap of effectors found in at least two haplotypes. Colors represent percent amino acid identity across rows. Haplotypes are shown to the right.

Next, we analyzed effector conservation across *C*Lso haplotypes. None of the identified effectors share over 44% amino acid identity with proteins outside of their genus, indicating that they are unique to Liberibacter and their function cannot be predicted by homology. Only one effector, HPE74, possessed an identifiable domain Cupredoxin_1 domain PF13473 (Table S1), which is predicted to bind a type I copper redox site and involved in electron transfer reactions (Dennison, 2013). To better understand the role of effector repertoires in the context of phylogenetic relationships, we performed phylogenetic analyses of *C*Lso haplotypes using 825 orthologous genes and compared it with phylogeny generated from the 13 core effectors (Fig 1d). In general, the topology of both trees was similar. Analyses of variable and core effectors across haplotypes revealed that haplotypes A and B and haplotypes C and D are the most similar to each other (Fig 1c, 1e), which is incongruent with their phylogeny but does coincide with their host range (Figs 1d, e). We observed variation at the nucleotide level for the same effector between haplotypes (Fig 1e). For example, HPE2, which is a core effector, possesses 80-93% amino acid similarity between haplotypes. It is possible that the combination of genome reduction associated with intracellular lifestyle, high AT content, and continuing expansion of Liberibacter geographical range are driving these observations.

### 2.2 *C*Lso SEC-effectors localize to different host compartments

Haplotypes A and B possess a similar repertoire of effectors and exhibit similar host range, causing disease in potato and tomato (Fig 1b-e). Haplotype B is more aggressive on both tomato and potato (Wen *et al*., 2013). Therefore, we focused on SEC-dependent effectors shared between haplotypes A and B (Fig 1c) or unique for haplotype B.

To gain insight into the possible role of *C*Lso SEC-dependent effectors, we evaluated their subcellular localization by fusing the mature effector (without signal peptide) with an N-terminal turboGFP (tGFP) tag using Golden Gate technology (Fig 2a, Fig S1). turboGFP is a copGFP variant with more rapid folding and increased fluorescence (Shagin *et al*., 2004). tGFP-mEffector fusions were visualized by confocal microscopy using *Agrobacterium-*mediated transient expression in *Nicotiana benthamiana.* Effector expression was verified using anti-tGFP immunoblot (Fig S2). We were able to detect full-length protein expression for the majority of effectors tested (Fig S2). We were unable to detect GFP expression for five effectors using anti-tGFP immunoblot but were able to visualize them by confocal microscopy (HPE2, HPE7, HPE16, HPE21 and HPE73) (Fig S1-2, data not shown).

**Figure 2.**
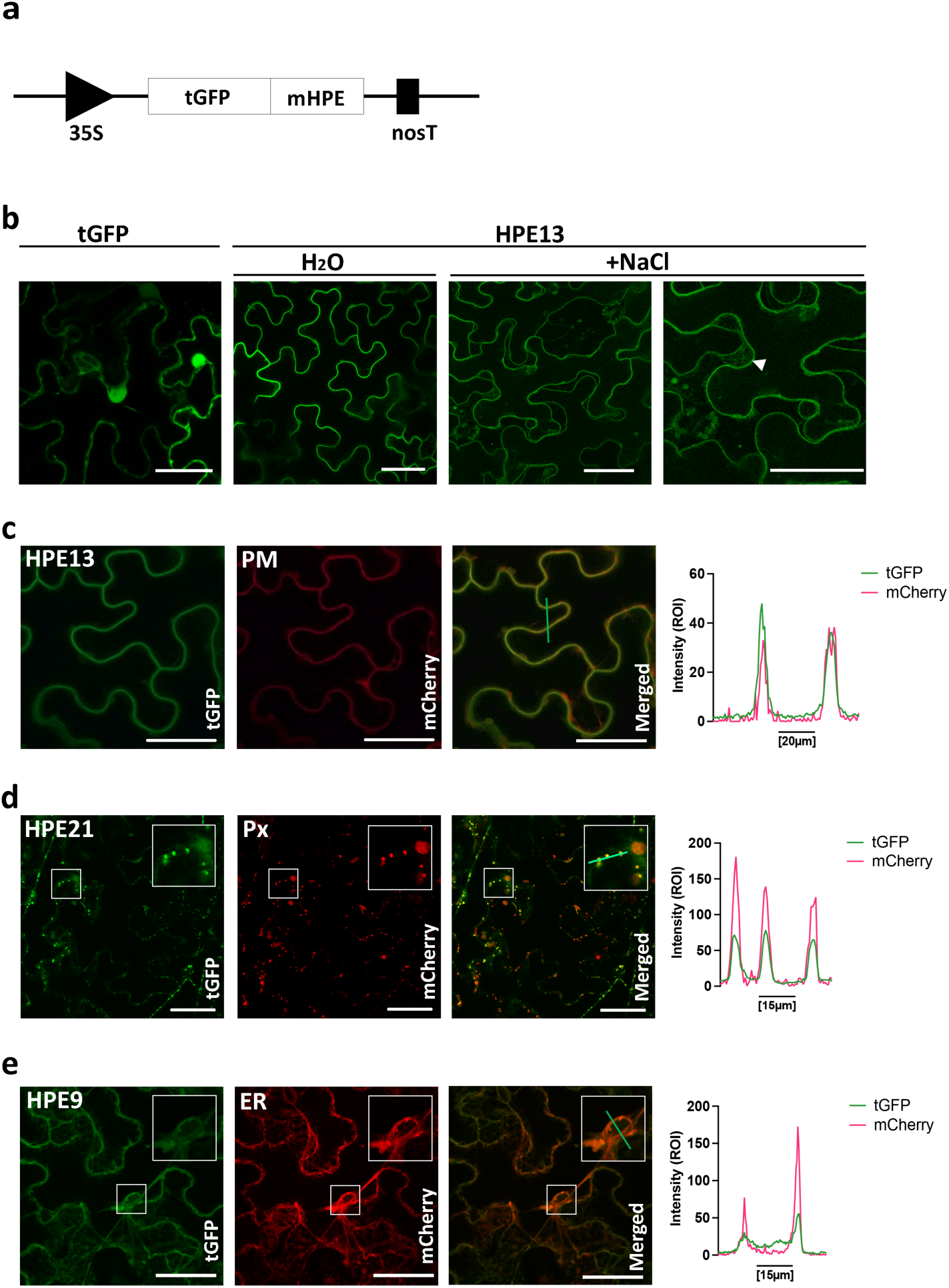
*C*Lso effectors localize to specific plant subcellular compartments. **a.** Mature effectors (mEffector) lacking their N-terminal signal peptide were cloned with an N-terminal fusion to TurboGFP (tGFP) and visualized by confocal microscopy 24h after transient expression in *N. benthamiana.* **b.** tGFP-HPE13 localizes to the plasma membrane. Leaves expressing HPE13 were subjected to plasmolysis with 1 M NaCl for 30 min, arrow indicated Hechtian strands. Leaves infiltrated with water were used as a control **c.** tGFP-HPE13(left) was co-infiltrated with *AtPIP2A*-mCherry (center). Right panel shows the intensity profile for the green line in the merged image. **d.** tGFP-HPE21(left) was co-infiltrated with 1-PTS1 peroxisomal-targeted mCherry (center). Inset in the right panel is a close-up of the colocalization of tGFP-HPE21 with peroxisomes in the merged image. Right panel shows the intensity profile for the green line in the merged image. **Ee.** tGFP-HPE9 was co-expressed with a SP-mCherry-HDEL labeling the ER. Inset in the right panel is a close-up of the colocalization of tGFP-HPE9 with the ER in the merged image. Right panel shows the intensity profile for the green line in the merged image. Bars= 50µm.

HPE13 and HPE21 are present in a genomic island that is unique to haplotype B and exhibit different subcellular localizations (Table S1). We observed HPE13 in the plasma membrane (Fig 2b). To confirm HPE13 localization, we induced plasmolysis with 1M NaCl and were able to observe Hechtian strands stretching between the plasma membrane and the cell wall in tGFP-HPE13 expressing cells (Fig 2b, right panel). To further confirm the localization of HPE13, we co-expressed tGFP-HPE13 with the plasma membrane aquaporin marker *AtPIP2A*-mCherry. To confirm the colocalization of HPE13, we plotted the intensity profiles of each fluorophore across a linear section and overlapping fluorescence intensity profiles confirmed the colocalization of HPE13 with the PM marker (Figure 2c). Bacteria occasionally hijack the palmitoylation machinery of the host cell and undergo lipidation to accumulate in membranes. HPE13 was predicted to have a S-palmoytilation site in the C-terminus by GPS-Palm (Ning *et al*., 2021). This post-translational modification may target HPE13 to plant membranes.

In contrast to HPE13, HPE21 targets a eukaryotic organelle. HPE21 localized to punctate structures that were ubiquitously distributed in the cytoplasm. HPE21 did not co-localize with a Golgi marker (soybean a-1,2-mannosidase I) (Fig S3). Instead HPE21 targeted peroxisomes as indicated by co-localization with 1-PTS1 peroxisomal-targeted mCherry (Fig 2d). To confirm the colocalization of HPE21, we plotted the intensity profiles of each fluorophore across a linear section, as shown in Fig 2d. The intensity profiles of tGFP-HPE21 and the peroxisome marker matched exactly. HPE9 accumulated in the perinuclear membrane, a known site of ER accumulation (Perkins & Allan, 2021), where the ER marker SP-mCherry-HDEL also accumulates (Fig 2e).

tGFP-HPE9 mainly co-localizes with SP-mCherry-HDEL throughout the cell, although there are some differences in the patterns of each protein. For example, SP-mCherry-HDEL is in the ER lumen while HPE21 associates with the cytosolic face of the ER membrane. Altogether, these results suggest *C*Lso effectors target specific eukaryotic subcellular compartments.

### 2.3 *C*Lso effectors target specific subnuclear compartments

Multiple Phytoplasma effectors target plant nuclei to reprogram their hosts (Tomkins *et al*., 2018). We were keen to understand if nuclear targeting is a common strategy amongst phloem-limited plant pathogenic bacteria. Seven proteins were to encode a monopartite or bipartite nuclear localization signal (NLS): HPE2, HPE8, HPE9, HPE15, HPE16, HPE22 and HPE30 (Fig 3, Fig S1). However, only four effectors exhibited exclusive nuclear localization: HPE16, HPE18, HPE19 and HPE73, (Fig 3c, Fig S1). HPE18 is found in haplotypes A, B, and D; HPE19 is found exclusively in haplotypes B and D; HPE8, HPE16, and HPE73 are core effectors (Table S1). HPE8 preferentially accumulates in the nucleus but is also found in the cytoplasm (Fig 3b). Interestingly, we found HPE16 present in fast-moving punctate bodies inside the nucleus (Fig 3c, Video S1). These structures, known as nuclear speckles, are important for RNA metabolism and possibly facilitate regulation of gene expression (Reddy *et al*., 2012; Bazin *et al*., 2018). HPE18 accumulates in the nucleolus, while HPE73 accumulates in both the nucleus and nucleolus (Fig S1b). The majority of *C*Lso effector proteins, 16 out of 23 tested effectors, exhibited nuclear cytoplasmic localization (Fig 3, Fig 4, Fig S1). Five effectors exhibited sole or predominant nuclear localization. These data highlight the importance of effector nuclear targeting in Liberibacter.

**Figure 3.**
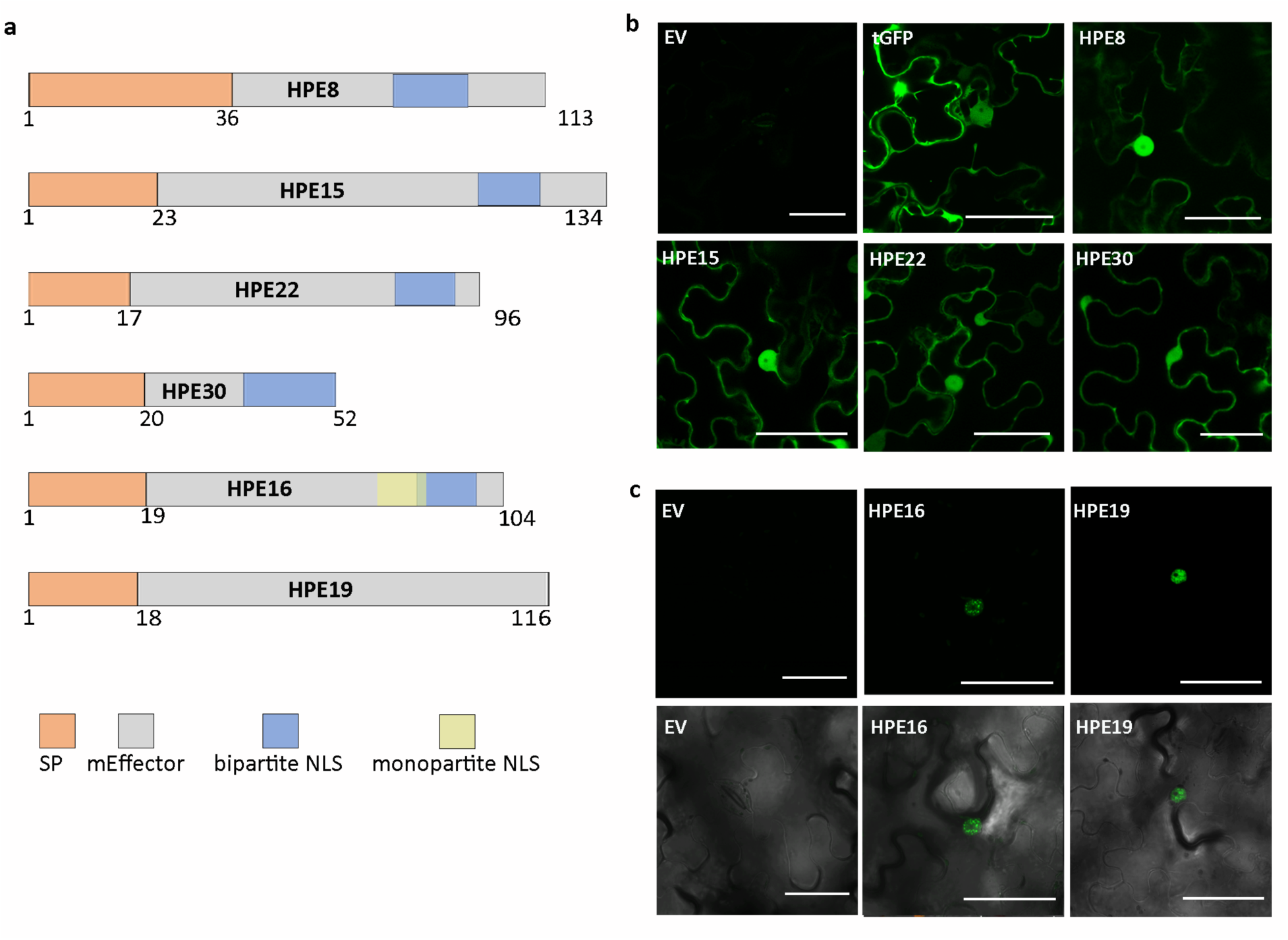
Multiple *C*Lso effectors localize to the nucleus and cytoplasm *in planta*. Mature effectors (mEffector) lacking their N-terminal signal peptide were cloned with an N-terminal fusion to TurboGFP (tGFP) and visualized by confocal microscopy 24h after transient expression in *N. benthamiana.* **a.** Schematic representation of *C*Lso-effectors including predicted nuclear localization signals (NLS). **b.** The majority of *C*Lso effectors exhibited a nuclear and cytoplasmic localization (16 out of 23 tested, see Fig S1). **c.** HPE16 and HPE19 exhibit nuclear localization, bottom panel merged with brightfield. Bars: 50µm

**Figure 4.**
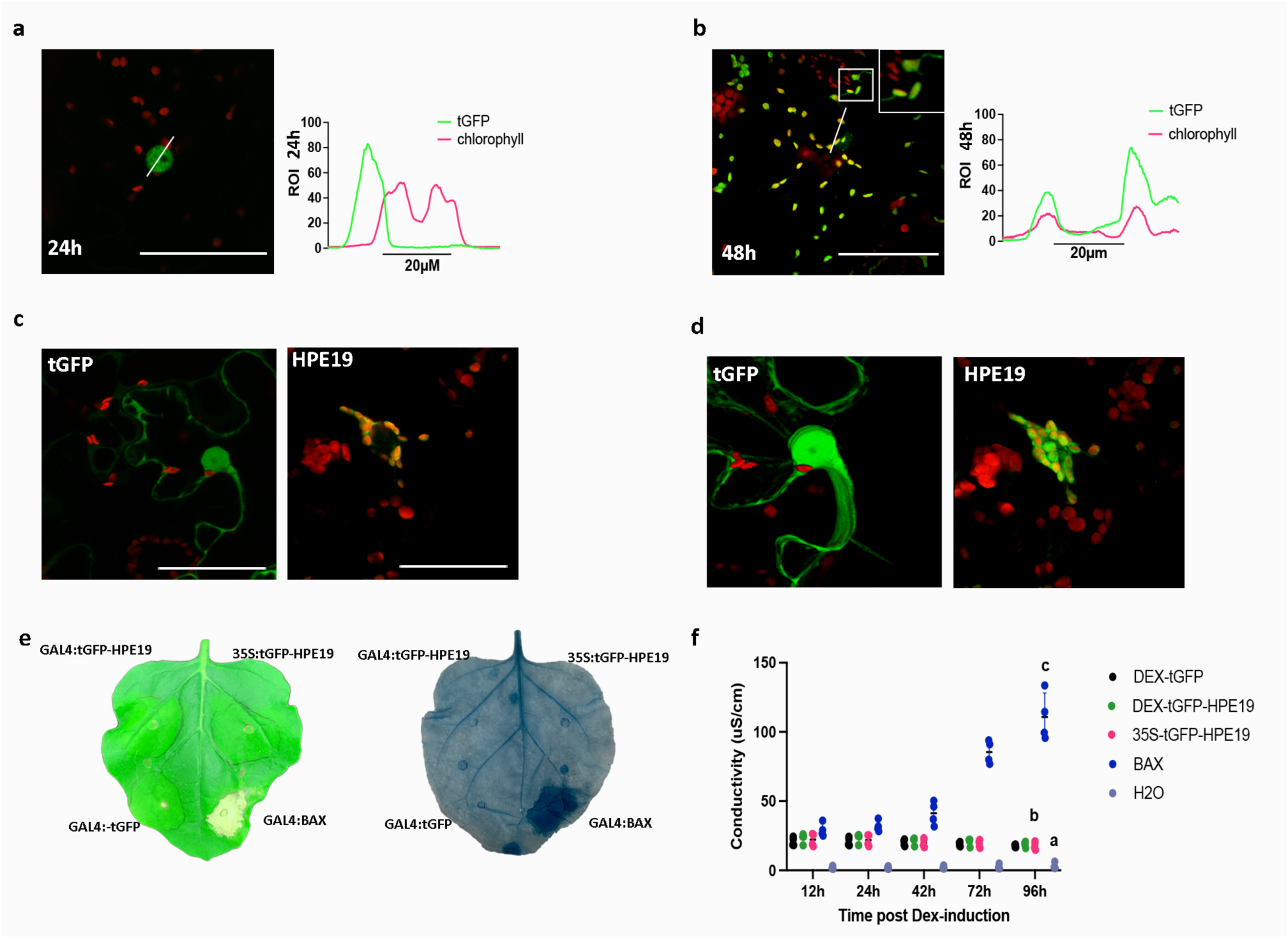
HPE19 exhibits dynamic nuclear chloroplast localization and alters immune responses in *Nicotiana benthamiana*. a-d. HPE19 lacking its signal peptide was cloned with an N-terminal fusion to TurboGFP (tGFP) under control of the GAL4 dexamethasone (Dex) inducible promoter, transiently expressed in *N. benthamiana*, and visualized by confocal microscopy. Dex was applied 12hpi **a.** HPE19 exhibits nuclear expression at 24hpi. Left panel shows GFP fluorescence, central panel includes chloroplast autofluorescence and right panel shows the intensity profile for tGFP and chlorophyll in the merged image. **b**. HPE19 primarily exhibits chloroplast localization at 48hpi. Insert in the central panel shows chloroplast stromule-like projections. Right panel, tGFP and chlorophyll intensity profiles overlap in the merged image. **c.** Representative image of nuclei expressing tGFP (left) and chloroplasts surrounding nuclei carrying tGFP-HPE19 (right). **d.** 3D reconstructions of the tGFP control (left) and tGFP-HPE19 (right) 48hpi illustrate chloroplast clustering around nuclei. 3D reconstructions were generated from 20 images. **e.** Overexpression of HPE19 fails to induce cell death Left: Macroscopic cell-death observed 5 days post-induction with 2uM Dex; Right: Microscopic cCell-death visualized after trypan blue staining observed 5 days post-induction with 2uM Dex **f.** Electrolyte leakage 12 to 96hpi after treatment with 2uM Dex. Individual dots represent values for four biological replicates per construct. Statistical differences were detected by a repeated measures ANOVA and different groups were detected using a Tukey multiple comparison test. Different letters indicate significantly different groups of means at 96hpi.

### 2.4 *C*Lso HPE19 targets nuclei and chloroplasts

Investigating HPE19 localization was challenging due to low-level expression. To enhance HPE19 expression, we cloned HPE19 with a tGFP tag under a dexamethasone inducible GAL4 promoter and induced expression with dexamethasone 12 hours post infiltration (hpi). tGFP-HPE19 expressed under an inducible promoter revealed differences in its subcellular localization overtime. At 24hpi, we observed nuclear localized HPE19 (Fig 4a). However, fluorescence was detected in primarily in chloroplasts and stromule-like projections 48hpi (Fig 4b, panel inset). HPE19 chloroplast localization was supported by overlapping tGFP fluorescence and chloroplast autofluorescence intensity profiles across a linear section at 48hpi but not 24hpi (Fig 4a-b right panel). Chloroplasts showing tGFP-HPE19 fluorescence were also observed nearby nuclei (4c, right panel). A 3D reconstruction of 20 images found that chloroplasts completely surround tGFP-HPE19 positive, but not control tGFP nuclei (Fig 4d). Clustering of chloroplasts around nuclei has been observed in plant-pathogen interactions and is believed to be important for plant defense activation (Krenz *et al*., 2012; Park *et al*., 2018a; Ding *et al*., 2019). Next, we investigated if expression of HPE19 induces plant cell death, a common defense response. Expression of HPE19 under a 35S or GAL4 inducible promoter did not induce a visible cell-death (Fig 4e). Trypan Blue vital staining revealed expression of HPE19 in *N. benthamiana* does not induce microscopic cell death. To confirm these results, we measured electrolyte ion leakage and were not able to detect any increase in ion leakage, a hallmark of cell death, after expression of HPE19 (Fig 4f). Expression of the pro-apoptotic protein BAX was able to induce visible cell death, cell death stained by Trypan Blue and electrolyte leakage (Fig 4 e-f). However, HPE19 slightly enhanced ROS and Ca^2+^ production in response to chitin perception (Fig 6). Contrary to our expectations, HPE19 suppressed ROS production in response to flg22, indicating this effector does not universally enhance plant responses to all pathogen features (Fig 6). It is possible that tGFP-HPE19 weakly activates plant defense or alters other processes resulting in stress response.

### 2.5 *C*Lso SEC-dependent effectors are able to move cell-to-cell

*C*Lso is a phloem-limited bacterial pathogen and is confined to sieve elements (Secor *et al*., 2009). Sieve elements are metabolically inactive enucleated cells that heavily rely on companion cells for their function. The specific localization of *C*Lso effectors in eukaryotic compartments indicates these proteins can move to companion cells and other neighboring cell types to target host compartments. To determine the ability of *C*Lso effectors to move intercellularly, mature *C*Lso effectors were C-terminally fused to eGFP and co-expressed with nuclear tdTomato (Fig 5a). As a positive control, we co-expressed the highly mobile eGFP with nuclear tdTomato. As a negative control, we co-expressed the nonmobile 2xeGFP with nuclear tdTomato. NLS-tdTomato and the 2xeGFP control are too large to diffuse cell-to-cell. A low concentration of *Agrobacterium* was used to facilitate single-cell transformation. Effector mobility was observed 24h after *Agrobacterium*-mediated transient expression in epidermal cells of *N. benthamiana*. Cell-to-cell movement was visualized in cells with green fluorescence surrounding an original transformation event. A clear positive result for this assay requires robust effector expression.

**Figure 5.**
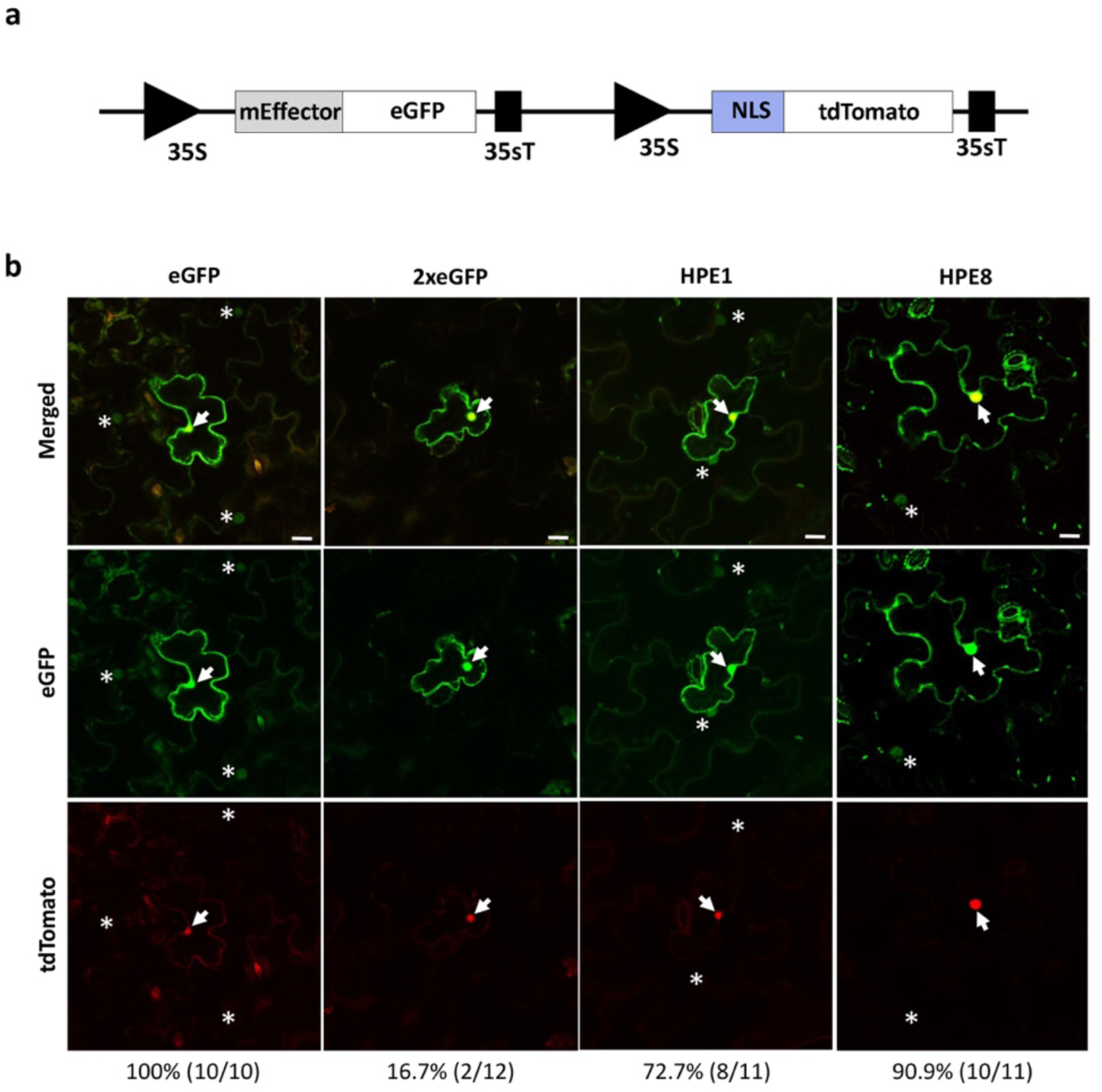
*C*Lso effectors are capable of cell-to-cell movement. **a.** Schematic representation of the construct used to test the ability of *C*Lso effectors to move cell-to-cell. Mature effectors (mEffector) lacking their N-terminal signal peptide cloned with a C-terminal fusion to eGFP in a binary vector also containing tdTomato targeted to the nucleus. **b.** Confocal images show the diffusion of *C*Lso mEffector-GFP fusion proteins in *N. benthamiana* epidermal cells 24h after transient expression. The concentration of *Agrobacterium* for transient expression was OD_600_= 0.0005, which enabled single cell transformation. The originally transformed plant cell exhibits both strong red fluorescence in the nucleus (arrows) and green fluorescencet signals. Effector movement is determined by the detection of GFP but not tdTomato signal in cells surrounding the transformed cell, indicated as asterisks. Scale bars = 20 µm. Numbers at the bottom indicate positive movement events out of total events observed.

**Figure 6.**
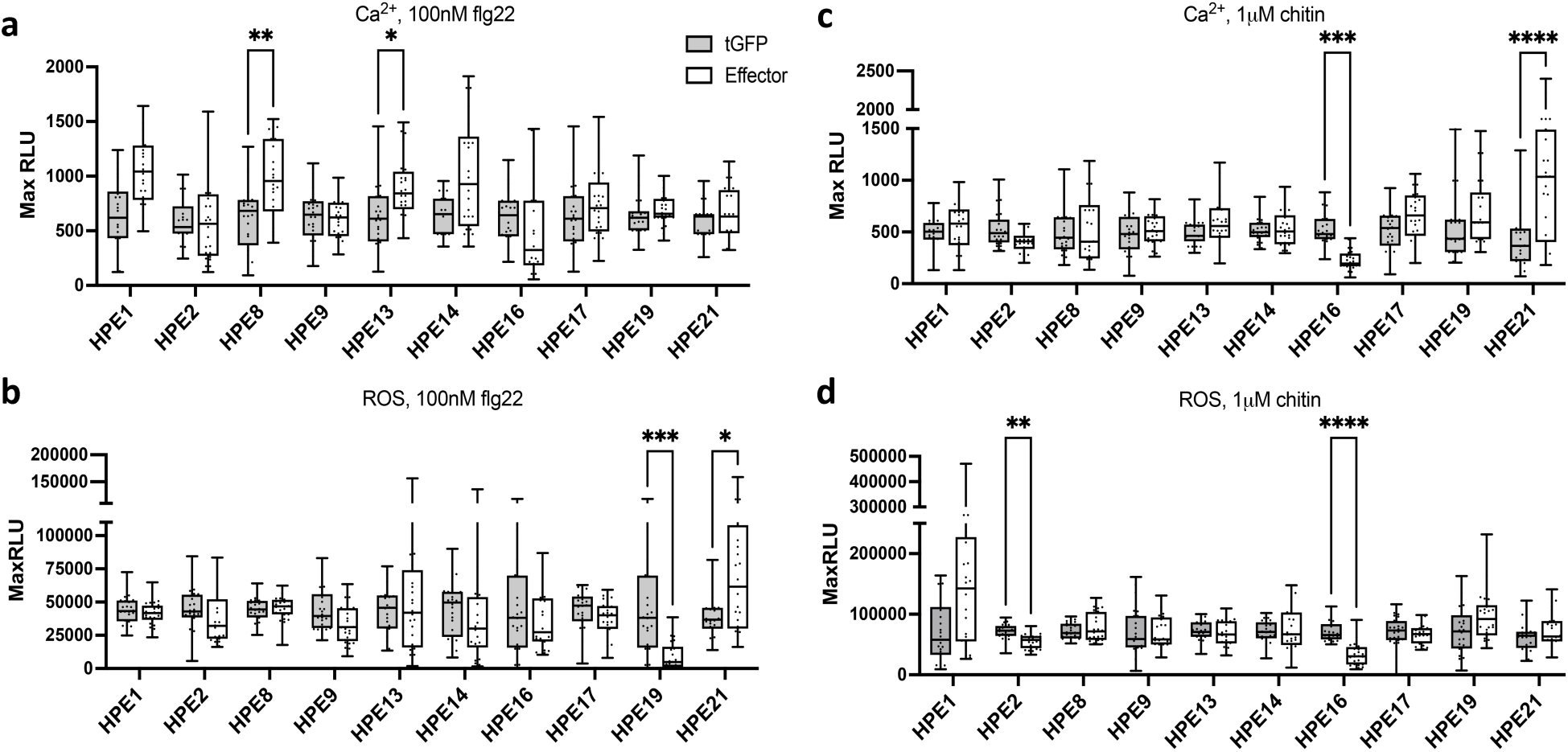
The majority of *C*Lso effectors do not alter plant immune responses to microbial features. Individual effectors were transiently expressed in *N. benthamiana* or the *N. benthamiana* Ca^2+^ reporter line SRLJ15 by *Agrobacterium* transient expression. *Agrobacterium* containing effector constructs were syringe infiltrated on a single leaf side-by-side with the tGFP control. Inoculations were performed by triplicate. Samples were taken 24h post-infiltration, floated in water or Coelenterezine solution for 16h and challenged with 100nM flg22 or 1µM chitin. ROS production and Ca^2+^ accumulation were measured on a luminometer. The majority of *C*Lso Sec-dependent effectors do not alter or weakly enhance the calcium cytosolic accumulation triggered by **a.** flg22 or **c.** chitin perception. The majority of *C*Lso Sec-dependent effectors do not suppress the ROS production elicited by **b.** flg22 or **d.** chitin. Statistical differences were detected by multiple Mann-Whitney tests. Multiplicity adjusted P values from the Holm-Sidak method were used to compute adjusted P values. *p<0.05, **p<0.005, ***p<0.0005, **\*\*\*\*** p<0.00005.

We were able to detect movement of two out of five tested *C*Lso effectors (HPE1, HPE8, HPE9, HPE16, HPE19). Out of 11 isolated single cell transformation events, we observed eight (72.7%) instances of HPE1 movement to adjacent cells (Fig 5b, S4). HPE1 was previously characterized as a plant cell death suppressor (Levy *et al*., 2019). We also observed movement of HPE8, which accumulates highly in the nucleus and is easy to visualize. Out of 11 isolated single cell transformation events, we observed 10 (90.9%) instances of HPE8 movement to adjacent cells (Fig 5b, S4). Differences in the size of HPE1 (11.32KDa) and HPE8 (8.78KDa) could account for the more limited movement of HPE1. Only 16.7% positives were observed for the 2xeGFP negative control, and 100% positives were observed for the eGFP positive control, indicating that the movement system is working properly. These data provide evidence that a subset of *C*Lso effectors are capable of cell-to-cell movement.

### 2.6 The majority of *C*Lso SEC-dependent effectors do not suppress plant immunity

The most well characterized function of pathogen effectors is their ability to suppress defense responses (Toruño *et al*., 2016). Insect feeding not only elicits wound associated responses, but their saliva and honeydew also carries insect and microbial features associated with plant pathogens and symbionts that can trigger a defense response (Huang *et al*., 2020). We investigated the ability of *C*Lso effectors to suppress plant immune defenses. In order to investigate immune suppression, we analyzed two early defense hallmarks of plant immunity: Ca^2+^ influx and production of ROS. These two responses also comprise the main events shaping subsequent defense responses reported in phloem (Gaupels *et al*., 2016, 2017).

We evaluated effector-mediated suppression of plant immune responses to two pathogen features: flg22 and chitin. Flg22 is an immunogenic peptide of flagellin perceived by the surface-localized receptor FLS2. Chitin is an oligosaccharide present in the cell walls of fungi and exoskeletons of insects that is perceived by LysM domain containing surface-localized immune receptors. Cytosolic Ca^2+^ accumulation was quantified in a transgenic *N. benthamiana* Aequorin reporter line (Segonzac *et al*., 2011). *C*Lso effectors were transiently expressed alongside with tGFP in *N. benthamiana*, 24h later, leaf tissue was collected and floated on coelenterezine solution for at least 12h and challenged with chitin or flg22. We analyzed ten effectors representing different subcellular localizations (Fig 3, 4, 6, S1). Surprisingly, *C*Lso effectors failed to suppress the intracellular accumulation of Ca^2+^ upon perception of flg22 or chitin (Fig 6a, 6c). Only HPE16 was able to inhibit the Ca^2+^ influx after chitin challenge. Several effectors increased Ca^2+^ influx upon perception of either flg22 (HPE8, HPE13) or chitin (HPE21) (Fig 6a, 6c). While some effectors displayed a similar trend of increased Ca^2+^ influx, these results were not statistically significant.

Next, we examined whether *C*Lso effectors alter ROS production in response to perception of flg22 and chitin. *C*Lso effectors were infiltrated alongside with tGFP and transiently expressed in *N. benthamiana*, 24h later leaf tissue was collected, floated in water overnight and challenged with either immunogenic feature. In general, the majority of tested effectors failed to suppress the ROS production elicited by either flg22 (Fig 6b) or chitin (Fig 6d). Only three effectors consistently inhibited ROS production (Fig 6b, 6d). HPE16 suppresses chitin induced ROS and Ca^2+^ influx, but does not significantly alter flg22-induced responses (Fig 6). HPE19 was able to suppress flg22-induced ROS, but slightly enhanced ROS and Ca^2+^ influx after chitin treatment (Fig 6). Differential suppression of flg22 and chitin immune outcomes have been reported for *Pseudomonas syringae* Type III effectors (Gimenez-Ibanez *et al*., 2018). *C*Lso effectors involved in plant immune suppression may target different immune signaling components. Altogether these data indicate that the majority of *C*Lso effectors are not capable of suppressing plant immune hallmarks. This pattern is strikingly different compared to the extensive defense suppressing effector activity of other Gram-negative foliar pathogens, including *P. syringae* (Guo *et al*., 2009).

### 2.7 Differential expression of *C*Lso effectors in the plant and the vector

Insect-vectored plant pathogens must be able to adapt to two drastically different environments: their vector and host. *C*Lso SEC-dependent effectors likely contribute to the ability to colonize plant sieve elements and propagate in *B. cockerelli*. We hypothesized that *C*Lso expresses different suites of effectors in vector and host, which can provide insight into contribution of specific effectors in disease development. We investigated the expression of *C*Lso effectors in tomato and in *B. cockerelli* using one-step RT-qPCR. Synchronized *B. cockerelli* colonies infected with haplotype B were used to transmit *C*Lso onto tomato cv. MoneyMaker (Fig 7a). Psyllids were kept in a muslin bag and 72h after vector transmission, removed for RNA extraction. Midribs were extracted from the original inoculated leaf one-month post-inoculation. Upregulated and downregulated genes in each treatment were identified after normalizing to the *C*Lso housekeeping gene *GlnA*. We were able to identify 5 effectors that are strongly expressed in tomato one-month post-inoculation compared to the psyllid (HPE2, HPE3, HPE16, HPE27, HPE74; Fig7b). We also identified 5 effectors that are strongly expressed in the psyllid compared to tomato (HPE4, HPE6, HPE15, HPE18, HPE33). Nine effectors were more uniformly expressed in both organisms (Fig7b). Collectively, these data support the hypothesis that some *C*Lso effectors are required for eukaryotic colonization, but other suites of effectors are differentially deployed to manipulate host and vector.

**Figure 7.**
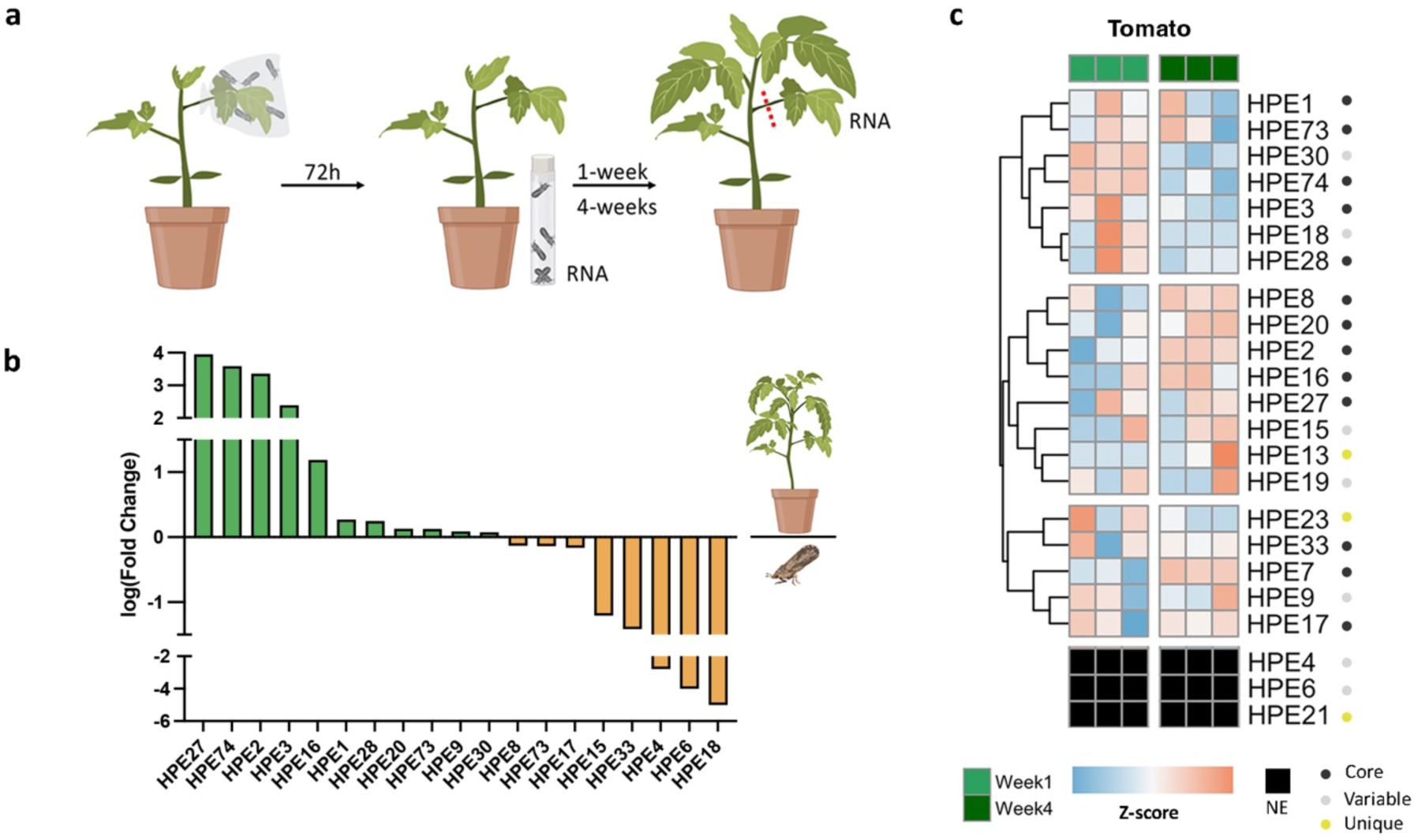
*C*Lso effectors exhibit dynamic expression according to organism and time. **a.** Experimental design for analyzing effector expression in psyllids (*B. cockerelli*) and plants (tomato). Fifteen one-day-old haplotype B psyllids were caged on the second leaf of four-week-old tomato cv Money Maker plants using a mesh bag. Seventy-two hours later, psyllids were removed and stored at -80°C for RNA extraction. One or four weeks later, midrib tissue from the originally infected leaf was collected for each biological replicate (n=3) **b.** Comparison of *C*Lso core, variable and unique effectors expression in tomato and psyllids. *C*Lso *GlnA* was used for normalization. Samples of the same organism cluster together. The △△Ct method was used to analyze effector expression, with results shown on a Log2 scale. **c.** Effector expression changes over time. The heatmap shows one experiment with three biological replicates and two different timepoints (one week and four weeks). Sets of effectors with similar expression patterns are shown as distinct clusters. *C*Lso *GlnA* was used for normalization. The △Ct method was used to analyze effector expression, with results shown as a Z-score. The pheatmap package in R was used to perform hierarchical clustering and visualize the results. NE= not expressed.

In infected plants, Liberibacter distribution and titers and disease symptoms are patchy and variable from plant-to-plant (Li *et al*., 2009a,b). We also observed differences in *C*Lso titers between plant samples where Ct values of the *C*Lso housekeeping gene *RecA* at week one varied between 26.89-34.0 and at week four varied between 22.74-24.61. Liberibacter infection is characterized by an asymptomatic period followed by the development of disease. In addition, disease symptoms can take months to years to develop, depending on the plant host. Due to the long latent period for disease development, we hypothesize that *C*Lso differentially expresses effectors depending on infection stage. To this end, we also compared effector expression in tomato at one week and four weeks post-vector transmission.

Although there was variation between biological samples, we identified sets of early (Fig 7C, top panel) and late (middle panel) acting effectors. Interestingly, the majority of late acting effectors are core effectors (Fig 7c). In general, HPE74 and HPE30 exhibited reproducible expression and were highly upregulated at one week while HPE2 and HPE8 were reproducibly downregulated at four weeks post-vector transmission (Fig 7c). These results demonstrate that over the course of infection the effector expression profile shifts, potentially enabling adaptation to new environments.

## DISCUSSION

VBDs significantly impact agricultural production and are important emerging diseases. Despite the importance of VBDs, scientists do not have a robust understanding of host manipulation regulated by vector-borne pathogens compared to foliar pathogens (Perilla-Henao & Casteel, 2016; Huang *et al*., 2020). Some of the most important bacterial vector-borne pathogens reside in the Liberibacter genus, including the devastating citrus pathogen *C*Las and the Solanaceous and Apiaceous pathogen *C*Lso. In this study we analyzed the identity, localization, defense suppression and expression of *C*Lso haplotype A and B effectors. Our results show *C*Lso effectors target diverse eukaryotic subcellular compartments, are capable of moving cell-to-cell and exhibit complex expression patterns in vector and host. Most *C*Lso effectors do not suppress host immune responses, indicating that they have novel targets, potentially related to VBD spread.

Vector-borne plant pathogens must exhibit high transcriptional flexibility in order to adapt to distinct organisms, insect and plant. Bacterial effector expression is known to be hierarchical, but there is little evidence of dynamic expression patterns during the course of plant bacterial infection (Mills *et al*., 2008; Lara-Tejero *et al*., 2011; Portaliou *et al*., 2017). Filamentous pathogens are known to express waves of effectors during infection. For example, the hemibiotrophic fungal pathogen *Colletotrichum higginsianum,* expresses specific sets of effectors during pre-penetration, biotrophic and necrotrophic infection stages (Kleemann *et al*., 2012). Similarly, we observed *C*Lso effectors exhibit dynamic expression profiles in tomato, identifying sets of early and late expressed effectors (Fig 7c). The plant distribution and titer of *C*Lso is patchy (Li *et al*., 2009a). If *C*Lso effector expression is density dependent, this could explain the observed variability in effector expression across samples. Differences in effector expression were also observed between tomato and psyllid (Fig 7b), indicating that *C*Lso utilizes its effector repertoire to adapt to diverse environments. Several *C*Las effectors also show differential expression in host and vectors as well as different host genotypes, but dynamics during infection have not been studied (Yan *et al*., 2013; Pagliaccia *et al*., 2017; Clark *et al*., 2018; Liu *et al*., 2019). Our results suggest *C*Lso requires a specific set of effectors to interact with the plant at different stages of the infection and core effectors, which are predominantly expressed during late stages of infection, most likely play a role in disease development.

*C*Lso SEC-effectors target several subcellular compartments that are absent in bacteria including the ER, peroxisomes, chloroplasts and nuclei (Fig 2,3 and 4); these compartments are known targets of plant pathogen effectors (Toruño *et al*., 2016; Park *et al*., 2018b). SEC effector targeting of eukaryotic compartments indicates they function outside of the bacterium to modulate host and vector. In addition, we found *C*Lso effectors can move cell-to-cell (Fig 5), indicating they are capable of movement during infection. Effector mobility has been observed in other vascular (*Fusarium oxysporum* in tomato) and non-vascular (*Magnaporte oryzae* in rice) pathosystems (Khang *et al*., 2010; Cao *et al*., 2018). Movement of effectors might be particularly important for phloem-limited pathogens to thrive. Multiple Phytoplasma SEC-dependent effectors are mobile, target transcription factors and influence disease symptomology and transmission (Tomkins *et al*., 2018). *C*Lso HPE16 localizes to nuclear speckles (Fig 3c), a site of storage for RNA metabolism proteins, including transcription factors (Bazin *et al*., 2018). There are a few examples of effectors targeting this compartment, including PsAvh52, a *Phytophthora sojae* effector, that induces transcriptional reprogramming by recruiting a plant transacetylase to nuclear speckles (Li *et al*., 2018). Further characterization of the plant targets of mobile, nuclear localized effectors will reveal if targeting of transcription factors is a common mechanism for phloem-limited pathogens.

Plants rely on surface localized receptors to recognize pathogens including insects and are activated upon insect feed and salivary secretions in the apoplast. Phloem defense responses include the production of ROS, Ca^2+^ influx, callose deposition and activation of proteins capable of occluding sieve elements (Huang *et al*., 2020). However, a detailed mechanistic understanding of phloem-mediated defense remains elusive. Watery saliva of aphids and leafhoppers contain effectors that suppress plant recognition, demonstrating the importance of plant immunity against VBD (Huang *et al*., 2020). Transcriptional profiling in tomato has revealed that *C*Lso haplotype B infected plants exhibit downregulation of defense related genes during the initial stages of infection (Casteel *et al*., 2012; Huot *et al*., 2018). The nuclear effectors HPE16 and HPE19 as well as the nuclear-cytoplasmic effector HPE2 were successful at suppressing cytosolic Ca^2+^ accumulation and/or ROS production (Fig 6). *C*Lso HPE1 was previously reported as a variable BAX-induced cell death and Prf hypersensitive response suppressor (Levy *et al*., 2019). Plants infected with *C*Lso usually contains high amounts of ROS and other defense related compounds (Wallis *et al*., 2012; Kumar *et al*., 2017), the fact that we could not find more effectors that consistently suppress ROS production upon perception of pathogen features could partially explain the high accumulation of these compounds. The *C*Las SEC effector, SDE1, promotes plant defense suppression by targeting plant immune proteases in the phloem (Clark *et al*., 2018). *C*Las SDE15 is a broad defense suppressor that inhibits HR induced by *Xanthomonas citri* pv *citri* in citrus (Pang *et al*., 2020). Although multiple *C*Las SEC effectors have been investigated, only two (SDE1 and SDE15) have been demonstrated to suppress plant immunity (Clark *et al*., 2018; Pang *et al*., 2020). Similarly, in this study we found the majority of *C*Lso SEC dependent effectors fail to suppress early hallmarks of defense (Fig 6). These results suggest Liberibacter only requires a few effectors for defense suppression.

Extracellular bacteria must overcome a multilayered plant defense before being able to establish disease, therefore the majority of their effectors are involved in plant immunity suppression (Guo *et al*., 2009; Medina *et al*., 2018; Traore *et al*., 2019). In contrast, phloem-limited pathogens are deposited directly inside plant cells, escaping apoplastic defense. This protective intracellular niche could explain the low number of SEC-dependent effectors in Phytoplasmas and Liberibacters reported in plant defense suppression (Sugio et al., 2011b; Wang et al., 2018; Clark et al., 2018; Pang et al., 2020, this study). Other vector-borne pathogens manipulate their hosts for transmission. For example, the *Turnip mosaic virus* and *Potato virus Y* use NIaPro, a protease with dynamic vacuolar localization, to increase vector performance and suppresses aphid-induced callose deposition (Casteel *et al*., 2015; Bak *et al*., 2017). Because phloem limited pathogens cannot colonize a new host by themselves, it is likely *C*Lso effectors play a role in modifying their environment and facilitating transmission.

Here, we initiated the characterization of *C*Lso effectors. *C*Lso’s interaction with Solanaceous plants, including tomato and potato, is economically important and represents a more tractable system to investigate VBD and unravel novel phloem specific defense strategies. This study sets the stage for future mechanistic investigation of *C*Lso effector targets.

## AUTHOR CONTRIBUTIONS

RPA designed the research, performed research, analyzed data and wrote the paper; JZ performed effector mobility assays, AB generated constructs for effector mobility assays, SPT performed the phylogenetic analyses in Fig 1, CC helped design effector expression experiments, CF contributed new analytical tools for effector mobility assays, GC designed the research, analyzed data and wrote the paper. TT performed blind immune suppression assays with HPE16 and HPE2 to validate results. LPH performed initial *C*Lso effector predictions. All authors reviewed and approved the final version of the manuscript.

## DATA AVAILABILITY

Genome and effector sequences are available in NCBI using the accession numbers provided in Table S1. All other raw data are available upon request to GC.

