## Supplemental Figures 1-4 for "Effectors from a Bacterial Vector-Borne Pathogen Exhibit Diverse Subcellular Localization, Expression Profiles and Manipulation of Plant Defense"

**SUPPLEMENTAL MATERIALS** (4 Tables, 4 Figures, 1 video)

**Table S1. Predicted *C.* Liberibacter solanacearum effectors.** Effectors were identified using the pipeline in Figure 1. Name indicates the common name for each effector. Accession number in NCBI for each effector according to haplotype. Absence indicates that the effector is not present in that genome. Effector presence and absence was experimentally tested using PCR in haplotype A (HenneA) and B (ZC1).

**Table S2. Primers used for cloning in this study.** CDS1ns refers to the CDS1 no stop codon module in Golden Gate.

**Table S3. Plasmids and Golden Gate components used in this study.**

**Table S4. Primers used in this study to evaluate effector expression by qPCR.** NCBI ID refers to the nucleotide sequence.

**Figure S1. The majority of *C*Lso effectors exhibit nuclear-cytoplasmic localization in *Nicotiana benthamiana*.** Mature effectors lacking their N-terminal signal peptide were cloned with an N-terminal fusion to TurboGFP (tGFP) and visualized by confocal microscopy 24h after *Agrobacterium*-mediated transient expression in *N. benthamiana.* **a.** The majority of *C*Lso effectors exhibited a nuclear and cytoplasmatic localization (16 out of 23 tested, see Figure 3B). **b.** Nuclear localized effectors. HPE18 mainly localizes in the nucleolous. HPE73 exhibits nuclear and nucleolar localization. Scale bars: 30µm

**Figure S2. Expression of *C*Lso effectors expressed in *N. benthamiana.*** Individual effectors were cloned as TurboGFP (tGFP) fusions and expressed in *N. benthamiana* using *Agrobacterium*-mediated transient expression. Total protein was isolated 24h post-infiltration (hpi). Full-length proteins were detected by immunoblot using anti-tGFP. CBB= Coomasie brilliant blue staining. **a.** Expression of HPE9 (37,36KDa), HPE3 (31,34KDa), HPE4 (32,14 KDa), HPE15 (38,95 KDa), HPE6 (30,66 KDa), HPE1 (37,36 KDa), and HPE14 (30,77 KDa) **b.** Expression of HPE1 (37,36 KDa), HPE13 (39,38 KDa), HPE17 (33,95KDa), HPE19 (36,32 KDa), HPE21 (43,41 KDa), HPE22 (34,13 KDa), and HPE30 (29,61 KDa) **c.** Expression of HPE18 (36,55 KDa), HPE5 (33,87 KDa), and HPE3 (31,34 KDa) **d.** Expression of HPE18 (36,55 KDa), HPE33 (31,7 KDa), HPE73 (32,81 KDa) and HPE74 (32,63KDa). **e.** Expression of HPE19 (36,62 KDa) was enhanced using an inducible promoter system. 2µM of dexamethasone was infiltrated 12h before harvesting tissue. HPE16, HPE2, HPE7, and HPE20 (not shown) were visualized confocal microscopy but not detected by immunoblot.

**Figure S3. HPE21 does not localize to the golgi.** Mature HPE21 effector lacking its N-terminal signal peptide was cloned with an N-terminal fusion to TurboGFP (tGFP) and co-expressed with organelle tagged mCherry. Co-localization was visualized by confocal microscopy 24h after *Agrobacterium*-mediated transient expression in *Nicotiana benthamiana.* HPE21 localizes punctate structures (left). tGFP-HPE21 was co-expressed with the mCherry tagged Golgi maker (the Golgi marker (soybean a-1,2-

mannosidase I). tGFP-HPE21 do not colocalize with Golgi bodies. Scale bars = 50 μm, BF = bright field.

**Figure S4. Complete microscopy panels for visualization of Lso effector movement in Figure 5.** Lso effectors were visualized after single cell transformation events. Mature effectors lacking their N-terminal signal peptide were cloned with a C-terminal fusion to eGFP in a binary vector also containing tdTomato targeted to the nucleus. Effector movement was visualized by confocal microscopy 24h after *Agrobacterium*-mediated transient expression in *Nicotiana benthamiana.* The concentration of *Agrobacterium* for transient expression was $OD_{600}$= 0.005, which enabled single cell transformation. Panels correspond to entire image taken for each sample in Figure 5. The originally transformed plant cell exhibits both strong red fluorescence in the nucleus (arrows) and green fluorescent signals. Effector movement is determined by the detection of GFP but not tdTomato signal in cells surrounding the transformed cell, indicated as asterisks. Scale bars = 20 μm.

**Video S1. HPE16 is present in fast-moving punctate bodies in the nucleus.**

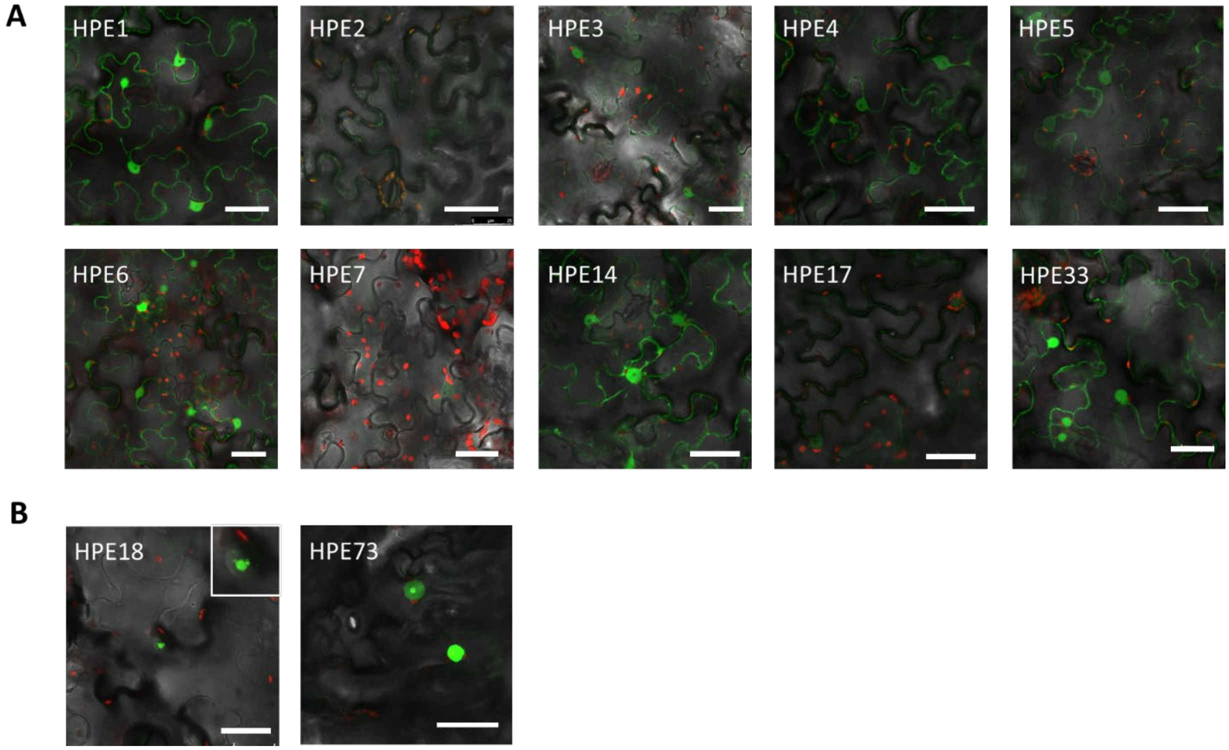


**Figure S1. The majority of Lso effectors exhibit nuclear-cytoplasmic localization in *Nicotiana benthamiana*.** Mature effectors lacking their N-terminal signal peptide were cloned with an N-terminal fusion to TurboGFP (tGFP) and visualized by confocal microscopy 24h after *Agrobacterium*-mediated transient expression in *N. benthamiana.* **a.** The majority of Lso effectors exhibited a nuclear and cytoplasmatic localization (16 out of 23 tested, see Figure 3B). **b.** Nuclear localized effectors. HPE18 mainly localizes in the nucleolous. HPE73 exhibits nuclear and nucleolar localization. Scale bars: 30µm

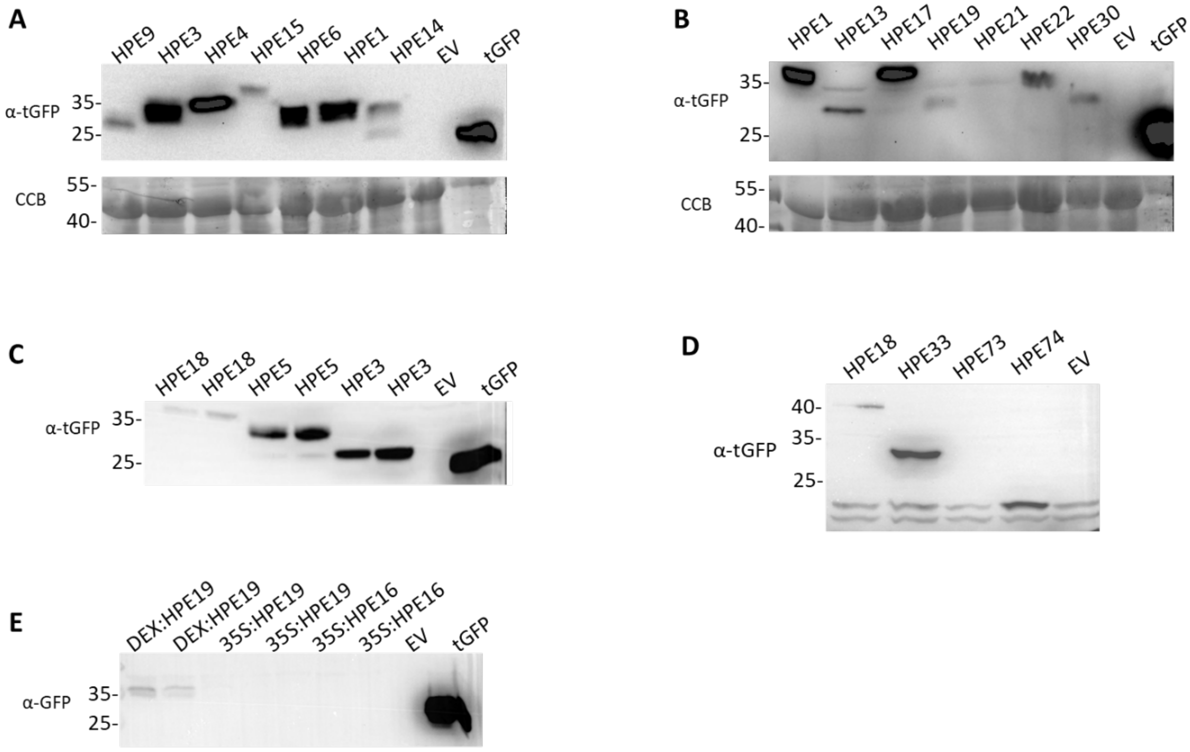


**Figure S2. Expression of Lso effectors expressed in *N. benthamiana.*** Individual effectors were cloned as TurboGFP (tGFP) fusions and expressed in *N. benthamiana* using *Agrobacterium*-mediated transient expression. Total protein was isolated 24h post-infiltration (hpi). Full-length proteins were detected by immunoblot using anti-tGFP. CBB= Coomasie brilliant blue staining. **a.** Expression of HPE9 (37,36KDa), HPE3 (31,34KDa), HPE4 (32,14 KDa), HPE15 (38,95 KDa), HPE6 (30,66 KDa), HPE1 (37,36 KDa), and HPE14 (30,77 KDa) **b.** Expression of HPE1 (37,36 KDa), HPE13 (39,38 KDa), HPE17 (33,95KDa), HPE19 (36,32 KDa), HPE21 (43,41 KDa), HPE22 (34,13 KDa), and HPE30 (29,61 KDa) **c.** Expression of HPE18 (36,55 KDa), HPE5 (33,87 KDa), and HPE3 (31,34 KDa) **d.** Expression of HPE18 (36,55 KDa), HPE33 (31,7 KDa), HPE73 (32,81 KDa) and HPE74 (32,63KDa). **e.** Expression of HPE19 (36,62 KDa) was enhanced using an inducible promoter system. 2μM of dexamethasone was infiltrated 12h before harvesting tissue. HPE16, HPE2, HPE7, and HPE20 (not shown) were visualized confocal microscopy but not detected by immunoblot.

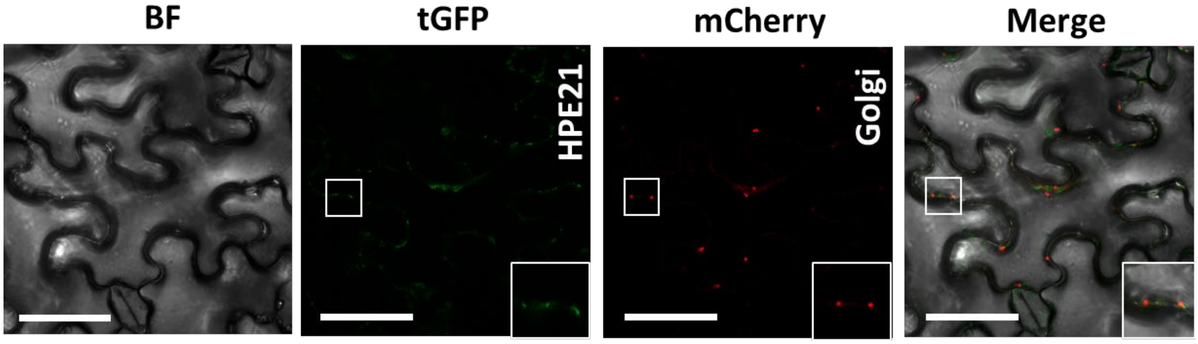


**Figure S3. HPE21 does not localize to the golgi.** Mature HPE21 effector lacking its N-terminal signal peptide was cloned with an N-terminal fusion to TurboGFP (tGFP) and co-expressed with organelle tagged mCherry. Co-localization was visualized by confocal microscopy 24h after *Agrobacterium*-mediated transient expression in *Nicotiana benthamiana.* HPE21 localizes punctate structures (left). tGFP-HPE21 was co-expressed with the mCherry tagged Golgi maker (the Golgi marker (soybean a-1,2-mannosidase I). tGFP-HPE21 do not colocalize with Golgi bodies. Scale bars = 50 μm, BF = bright field.

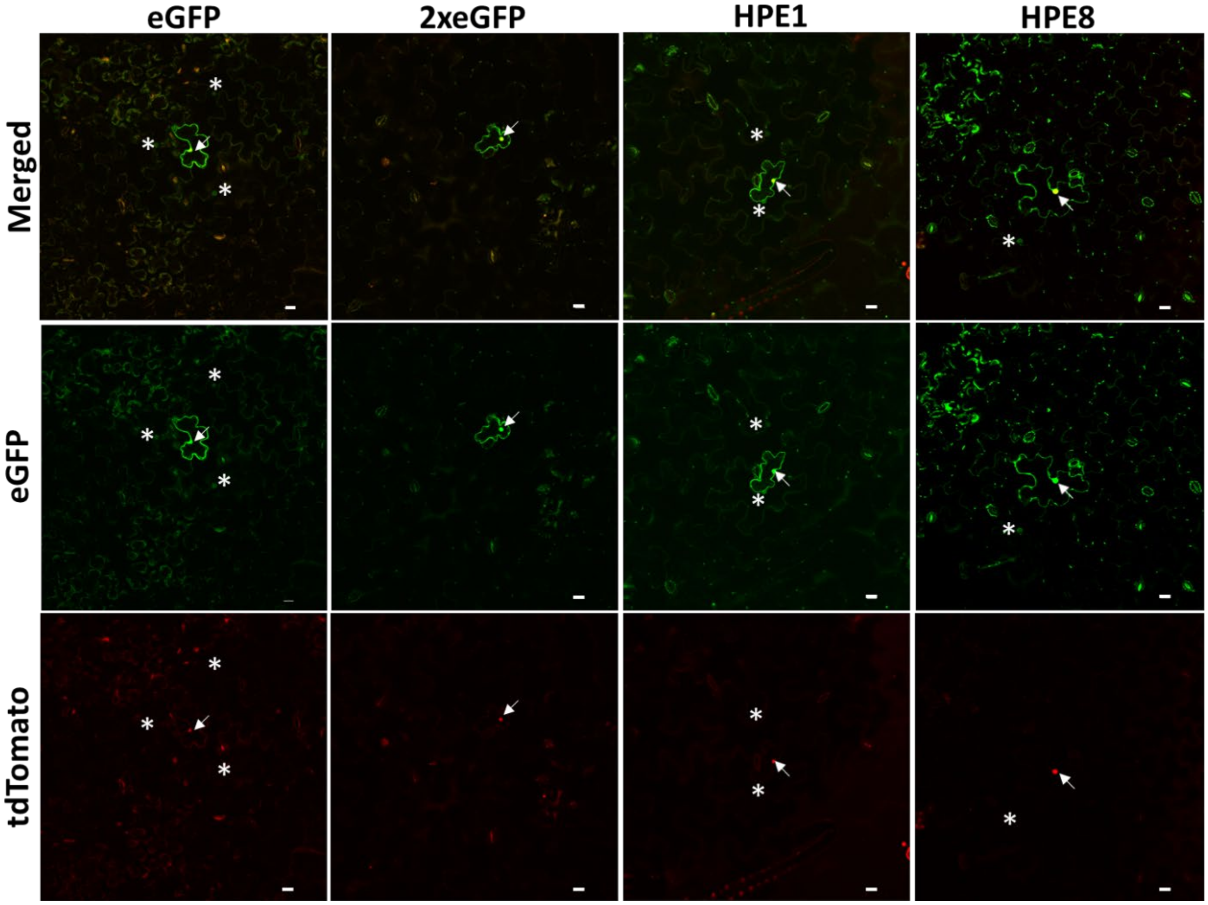


**Figure S4. Complete microscopy panels for visualization of Lso effector movement in Figure 5.** Lso effectors were visualized after single cell transformation events. Mature effectors lacking their N-terminal signal peptide were cloned with a C-terminal fusion to eGFP in a binary vector also containing tdTomato targeted to the nucleus. Effector movement was visualized by confocal microscopy 24h after *Agrobacterium*-mediated transient expression in *Nicotiana benthamiana.* The concentration of *Agrobacterium* for transient expression was $OD_{600}$= 0.005, which enabled single cell transformation. Panels correspond to entire image taken for each sample in Figure 5. The originally transformed plant cell exhibits both strong red fluorescence in the nucleus (arrows) and green fluorescent signals. Effector movement is determined by the detection of GFP but not tdTomato signal in cells surrounding the transformed cell, indicated as asterisks. Scale bars = 20 μm.
